# NMDA receptor ablation in medial prefrontal cortex disrupts value updating and reward history integration

**DOI:** 10.64898/2025.12.06.692679

**Authors:** Evan Knep, Angelica Velosa, Dana Mueller, Cathy Chen, Sophia Vinogradov, Matthew V. Chafee, Becket Ebitz, Sarah Heilbronner, Patrick E. Rothwell, Nicola Grissom

**Author notes:** to whom correspondence should be addressed: Nicola Grissom, Department of Psychology, University of Minnesota, Minneapolis, MN 75 East River Road, Minneapolis, MN 55454.

## Abstract

Schizophrenia, a serious mental illness, is associated with evidence of NMDA receptor (NMDAR) dysfunction and characterized by cognitive impairments that reflect impaired value updating and feedback-driven control; however the cellular and circuit-level mechanisms underlying these disruptions remain unclear. Here we test how NMDA receptor (NMDAR) signaling in the medial prefrontal cortex (mPFC) contributes to adaptive decision-making by combining targeted genetic ablation in mice and systemic pharmacology. Using a CRISPR-Cas9 approach to eliminate the obligate GluN1 subunit, we induced spatially confined NMDAR hypofunction in mPFC and compared its effects to systemic pharmacological blockade with the NMDAR antagonist MK-801 during performance of a touchscreen-based restless bandit task. Prefrontal NMDAR ablation impaired value discrimination, weakened the use of negative feedback, and reduced mutual information between recent outcomes and current choices, indicating disrupted reward-history integration. Reinforcement-learning models incorporating a choice-kernel term best captured behavior and revealed that NMDAR ablation selectively dampened learning and choice-history parameters governing flexible updating. Systemic MK-801 produced broad impairments in control animals, reducing accuracy, mutual information, and outcome sensitivity, yet exerted only modest additional effects after NMDAR ablation, suggesting that prefrontal NMDAR loss occluded much of the pharmacological disruption. Simulations using fitted RLCK parameters reproduced these patterns, showing convergent flattening of choice dynamics under MK-801 and persistent deficits in GluN1 ablated animals. Together, these findings demonstrate that prefrontal NMDAR signaling is necessary for effective value updating and feedback-driven learning, and that its loss recapitulates core features of systemic NMDAR hypofunction. This work establishes a mechanistic bridge between localized cortical glutamatergic dysfunction and the reinforcement-learning disturbances characteristic of schizophrenia.

## Introduction

Individuals with schizophrenia show disrupted reward-guided learning and difficulty maintaining optimal choice policies in dynamic environments, reflecting abnormalities in neural systems that support reinforcement learning and cognitive control [1–5]. These impairments most strongly predict real-world outcomes in patients, yet remain poorly understood mechanistically [6]. Translational task frameworks such as the multi-armed bandit paradigm have become powerful tools for bridging human and animal research on these processes. The bandit task requires continual adaptation to shifting reward contingencies, allowing quantification of latent components of decision making such as outcome sensitivity and value updating [7–13]. These tasks provide computationally precise measures that map onto conserved neural circuits of decision making, offering a way to test mechanistic links between behavior and underlying neurobiology relevant to schizophrenia.

One influential mechanistic account links cognitive impairments in schizophrenia to hypofunction of the N-methyl-D-aspartate receptor (NMDAR), which is essential for synaptic plasticity, long-term potentiation and depression, and higher-order cognition. Postmortem and molecular studies show reduced NMDAR subunit expression and altered maturation of prefrontal NMDAR function in individuals with the illness [14–18]. Systemic administration of agents that block NMDAR such as ketamine, phencyclidine, and MK-801 produce schizophrenia-like disruptions in cognition, perception, and neural synchrony across species [19–23]. NMDARs are highly expressed in regions that support value-based decision making and cognitive control, notably including the prefrontal cortex (PFC), hippocampus, and striatum [24–27]. The PFC has been a particular focus of research in schizophrenia because of longstanding evidence of alterations in both prefrontal activation and connectivity patterns in patients [28–30]. However, a major unanswered question is whether NMDAR hypofunction in PFC specifically is directly related to the cognitive impairments seen in schizophrenia.

To directly test how prefrontal NMDAR hypofunction contributes to schizophrenia-related cognitive impairments, we used a targeted genetic approach focused on the obligate GluN1 subunit of the receptor. Because GluN1 is required for assembly and function of all NMDAR, its loss effectively eliminates receptor activity within affected neurons, providing a precise means of inducing NMDAR hypofunction [31,32]. Using a CRISPR-Cas9 viral strategy, we selectively ablated GluN1 in the medial prefrontal cortex (mPFC), achieving a spatially precise reduction of NMDAR signaling while avoiding the widespread or developmental effects characteristic of systemic models. This model has been shown to produce early synaptic disruption followed by compensatory changes in excitatory and inhibitory transmission within mPFC circuits[33]. To contrast local with global disruption, we paired this manipulation with pharmacological NMDAR blockade using MK-801, predicting that prefrontal NMDAR ablation would reproduce key schizophrenia-like deficits in animals with intact GluN1 expression, but that MK-801 would produce fewer or minimized additional effects in animals with PFC GluN1 ablation consistent with a two-hit framework [34]. Cognition was assessed using a spatial variant of a two-armed restless bandit task implemented in a touchscreen operant chamber.

Selective prefrontal NMDAR ablation impaired value discrimination, reduced sensitivity to negative outcomes, and weakened integration of recent choice-outcome history, consistent with disrupted reinforcement learning dynamics. Systemic MK-801 produced widespread cognitive impairments in animals who had not undergone ablation, while animals with pre-existing prefrontal NMDAR ablation showed a reduced effect because these functions were already impaired. These results provide a direct comparison of local and systemic NMDAR disruption within a translational decision-making task framework, offering mechanistic insight into how prefrontal glutamatergic dysfunction may give rise to schizophrenia-related cognitive impairments.

## Methods

### Subjects

All procedures were approved by the University of Minnesota Institutional Animal Care and Use Committee (IACUC) and conducted in accordance with National Institutes of Health guidelines. Thirty-four mice (18 male, 16 female) expressing a Cre-dependent Cas9-GFP transgene[35] were bred in house and maintained on a C57BL/6J genetic background. The line is commercially available from Jackson Laboratories (Stock No. 026175). Animals were housed in temperature-controlled colony rooms (20.5°C; 69°F) on a 12h light-dark cycle (lights on 08:00–16:00). Behavioral testing was conducted during the dark phase. Mice had ad libitum access to water and were food restricted to maintain no lower than 85% of their free-feeding body weight. Testing occurred Monday-Friday, with free access to home cage chow provided on Fridays.

### Viral Vectors

Adeno-associated viral (AAV) constructs were produced by the University of Minnesota Viral Vector and Cloning Core. For selective disruption of the Grin1 gene, we utilized an AAV9 vector (AAV9-U6-gRNA[Grin1]-hSyn-mCherry-Cre; 2.16 × 10^13GC/ml) carrying a guide RNA sequence (CCGCGCCGATGTTGACAATCT) under the U6 promoter, and expressing an mCherry-Cre fusion protein driven by the human synapsin promoter, enabling both neuronal targeting and activation of Cre-dependent Cas9. Control mice were injected with a parallel AAV9 construct encoding a LacZ-targeted guide RNA (AAV9-U6-gRNA[LacZ]-hSyn-mCherry-Cre; 1.75 × 10^13GC/ml; guide sequence TGCGAATACGCCCACGCGAT). In both conditions, mCherry fluorescence was used to visualize virally transduced neurons. Procedures followed methods described in Dick et al., 2025[33].

### Surgical Procedures

Mice were approximately 22 weeks at the time of surgery. Animals were anesthetized with isoflurane gas delivered through a SomnoSuite system and positioned in a stereotaxic frame (Kopf Instruments, Tujunga, CA). Bilateral viral injections targeted the mPFC ( AP +1.80mm, ML ±0.35mm, DV −2.20mm from bregma). A total volume of 500nL was delivered into each hemisphere at 50nL/min over 10 minutes. The injection needle was left in place for an additional 10 minutes to facilitate diffusion before being slowly withdrawn. Animals received carprofen (10mg/kg, s.c.) on the day of surgery and for three consecutive days afterward.

### Drug Administration

To assess how NMDAR activity modulates bandit task strategy in Grin1 and control animals, we used MK-801 (dizocilpine), a non-competitive NMDAR antagonist. MK-801 was dissolved in sterile 0.9% saline, protected from light, and prepared fresh on each test day. Animals received intraperitoneal injections at 5ml/kg immediately before being placed in the chambers, which provided coverage for the ∼2-hour testing session[36]. On control days, animals received volume-matched saline. After a two-week saline-habituation period, mice completed three weeks of alternating MK-801 and saline sessions (Tuesday–Friday), yielding 12 test days. The MK-801 dose (e.g., 0.10mg/kg i.p.) was selected a priori based on prior work showing reliable effects on cognitive and decision processes in mice and rats [23,37–42].

### Behavioral Training

#### Restless Bandit Task

Following completion of the touchscreen acquisition protocol (see supplemental methods), mice were transitioned to the two-arm spatial restless bandit task. Each trial presented two identical visual stimuli on the left and right sides of the touchscreen (see supplemental methods for apparatus information). The reward probability associated with each side drifted independently, with a 10% chance of increasing or decreasing by 10% on each trial. Probabilities were constrained to avoid reaching absolute minima or maxima (i.e., never 0% or 100%).

Each session featured a novel, independently generated reward probability trajectory to prevent reliance on previous experiences. Mice completed bandit sessions Tuesday-Friday, with Mondays reserved for retraining on the 100–0 deterministic discrimination schedule. Sessions concluded after 300 trials or 120 minutes, and no testing occurred on weekends.

### Data Analysis

All analyses were conducted in R using custom scripts, with two-sided tests and α = 0.05 unless noted otherwise. Behavioral metrics included mutual information, negative outcome weight, and reinforcement-learning (RL) models that quantified trial-by-trial learning and choice dynamics. Mutual information assessed dependence of current choices on recent choice–outcome history, and negative outcome weight measured how losses influenced switching behavior. RL models (including Rescorla–Wagner, RL with choice kernels, stay-bias variants, and asymmetric learning-rate formulations) were fit separately for each animal and schedule using maximum-likelihood estimation with bounded L-BFGS-B optimization. Model selection relied on information-criterion comparisons. Full computational details—including model equations, parameter definitions, optimization procedures, and model comparison metrics—are provided in the Supplemental Methods.

## Results

### NMDAR Ablation Drives Decision Making Deficits

To test the role of prefrontal NMDAR function in value-based decision making, we examined how loss of the GluN1 subunit in the medial prefrontal cortex (mPFC) affected performance in a two-armed restless bandit task (**Figure 1A–B**). This task requires animals to make decisions in a challenging and highly dynamic environment. Animals chose between two locations whose reward probabilities drifted independently, requiring continuous tracking of changing contingencies and flexible updating based on recent outcomes. Before surgery, task performance did not differ between virus groups (Wilcoxon rank-sum test, *p* = 0.65) (Figure 1C). After mPFC GluN1 ablation, Grin1 animals showed reduced reward acquisition compared with LacZ controls (Wilcoxon rank-sum test, *p* = 0.0086). Within-group tests showed a trending but insignificant change in LacZ animals (Wilcoxon signed-rank test, p = 0.094) but a clear decline in Grin1 animals (Wilcoxon signed-rank test, p = 0.0012) (**Figure 1C**). Given that animals in this task typically operate only a few percentage points above chance, the observed group difference reflects a substantial shift in how effectively choices track the evolving reward structure across roughly ten thousand trials per animal (M = 9819 trials, SD = 877).

**Figure 1.**
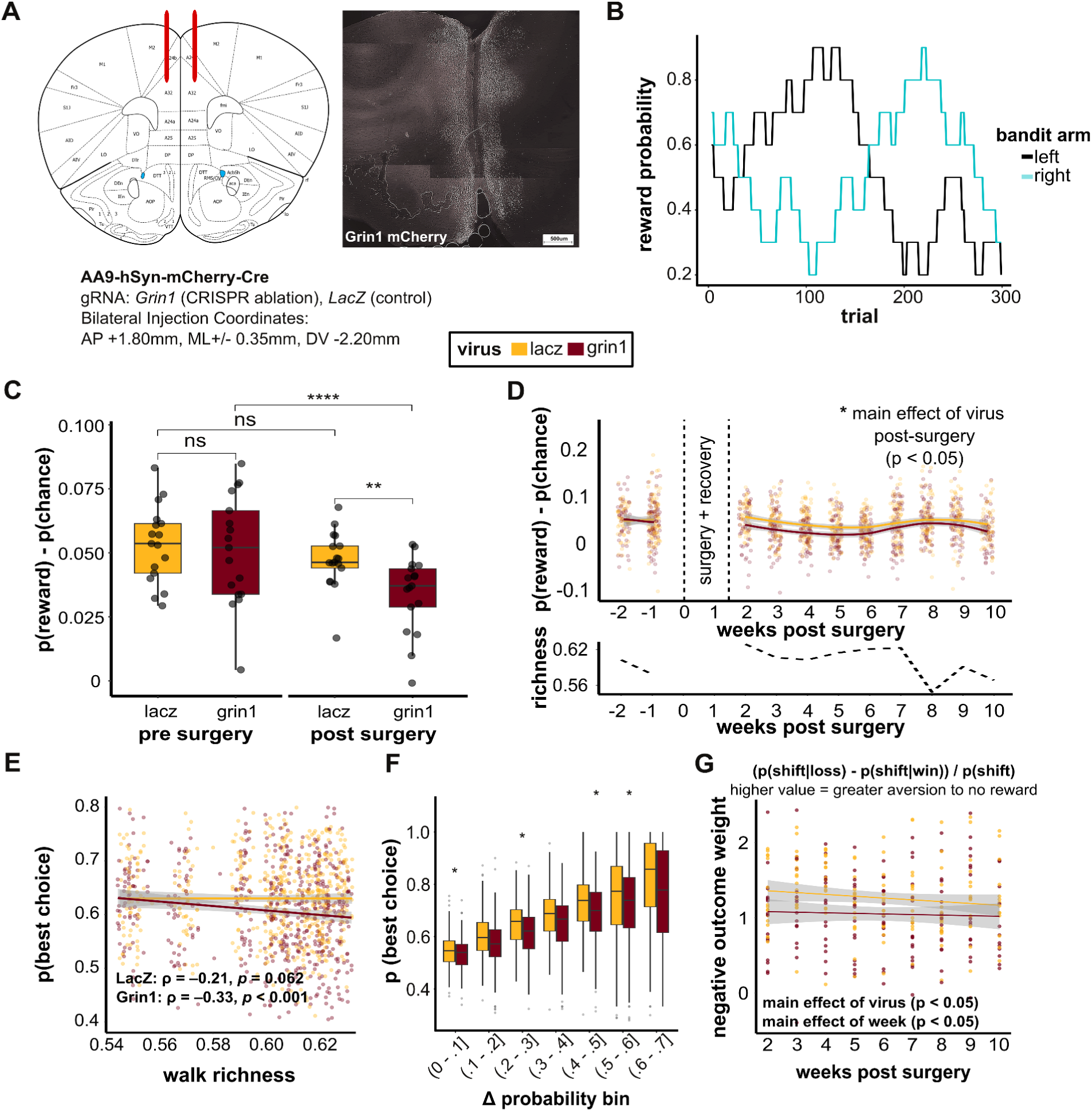
Prefrontal NMDAR ablation impairs performance and decision optimality. (**A**) Bilateral viral injection site within mPFC and construct identifiers (left) and representative image of viral-expression from grin1 ablation animal post experiment (right). (**B**) Example restless bandit walk showing changing reward probabilities for two options across 300 trials. (**C**) Reward minus chance compared between and within virus groups pre– and post-surgery. Within-group comparisons showed a trending but insignificant change in LacZ controls but a significant decrease in Grin1 animals. Post-surgery, Grin1 animals performed worse than controls, with no group difference pre-surgery. (**D**) Reward minus chance plotted over time and analyzed with a linear mixed-effects model (virus × weeks_post_surgery × richness, random intercept by animal). Grin1 animals performed worse overall than LacZ controls, and performance increased modestly with weeks post-surgery. The virus × week interaction showed a trend that was not significant, indicating that the group difference persisted across time with minimal compensation. Average walk richness (chance level) for each corresponding week plotted below. (**E**) Correlation between walk richness and accuracy assessed with Spearman correlations computed within animals (average across weeks 1–10). LacZ animals showed a modest negative trend, while Grin1 animals exhibited a stronger negative correlation, indicating that performance declined with increasing walk richness, particularly in Grin1 animals. (**F**) Accuracy across bins of Δprobability. A mixed-effects model revealed lower accuracy in Grin1 animals across the full Δprobability range, with no virus × bin interaction. Pairwise comparisons were consistent with this overall pattern, showing the largest group differences at low and intermediate Δprobabilities and numerically lower accuracy for Grin1 animals at higher bins. (**G**) Relationship between negative outcome weight and weeks post-surgery analyzed with a linear mixed-effects model (virus × week, random intercept by animal). Negative outcome weight decreased over time across animals, and Grin1 animals showed lower overall sensitivity to negative feedback than LacZ controls. Virus × week interaction was not significant.

Because the viral manipulation continues to express over time, we next examined how this group difference unfolded across the post-surgery period. A key feature of the restless bandit task is that its randomly drifting reward trajectories create day-to-day variability in the reward environment. This influences how useful outcomes are for learning, since richer walks, defined as sessions with a higher mean reward probability across trials, increase the likelihood that random choices are rewarded. To account for this variability across the duration of the experiment, the following analyses include walk richness. Performance across the eight post-surgery weeks showed a main effect of virus (β = –0.18, p = 0.032) and week (β = 0.02, p = 0.039), with no virus by week interaction (p = 0.09). This indicates a persistent deficit in the Grin1 group, and interestingly one that did not notably increase over the duration of viral expression and spread (**Figure 1D**). Richness significantly modulated weekly performance (week by richness interaction, p = 0.026), with a trend toward differential effects across virus groups (virus by richness interaction, p = 0.060), but importantly did not account for the observed group differences. A laboratory issue caused tissue degradation that prevented confirmation of viral expression in some animals, all primary observations above were replicated in the cohort of animals for which viral transfection could be confirmed, including significant group differences in p(reward) – p(chance) between Grin1 and LacZ groups, without a significant interaction between group and weeks post surgery (**Supplemental Figure 1**).

### NMDAR Ablation Reduces Decision Optimality and Increases Sensitivity to Outcome Noise

Week-to-week fluctuations suggested that group differences were shaped by how animals adapted to the changing reward structure, so we next assessed value discrimination independent of the overall reward rate by using the probability of selecting the “better” or more optimal option at any time (p(best choice)). In LacZ controls, optimal choice showed only a modest decrease with increasing walk richness (ρ = –0.21 ± 0.11, p = 0.062), whereas Grin1 animals showed a significant and more pronounced decline in best choices (ρ = –0.33 ± 0.06, p < 0.001) (**Figure 1E**). To determine whether differences in value discrimination contributed to this group effect, we analyzed p(best choice) across Δ-probability bins using a linear mixed-effects model. Grin1 animals showed lower selection of optimal choices overall than LacZ animals (main effect of virus, p = 0.036), indicating that the deficit was not restricted to low-evidence trials. Optimal choices increased systematically with larger Δ-probability values (main effect of bin, p < 0.001), and there was no virus × bin interaction (p = 0.56), showing that the groups were similarly influenced by value differences. Pairwise comparisons were consistent with this pattern, revealing the strongest group differences in low and intermediate Δ-probability bins, with numerically lower performance for Grin1 animals across the full range (**Figure 1F**).

### NMDAR Ablation Disrupts Feedback Sensitivity and Integration of Past Experience

We examined overall shift behavior (*p*Shift), defined as the probability of switching to the alternative option on the following trial. Grin1 and LacZ animals showed comparable average rates of shifting, with no significant group-level differences (*p* > 0.5). Overall shift rate alone does not capture how feedback guides behavior. To address this, we analyzed a normalized measure of negative outcome weight, which quantifies how strongly outcomes influenced shifting within each animal. This measure reflects the extent to which animals adjust their choices based on reward feedback, calculated as the difference in shift probability following a loss versus a win, normalized by overall shift rate. A linear mixed-effects model revealed a clear group difference: Grin1 animals displayed significantly lower negative outcome weight relative to controls (β = –0.35, *p* = 0.042) (**Figure 1G**). This reduction suggests that Grin1 animals were more willing to persevere through unrewarded trials rather than adjusting their behavior. The model also identified a main effect of week (β = –0.038, *p* < 0.001), indicating that both groups gradually decreased their negative outcome weight across weeks, with no significant virus × week interaction (*p* = 0.12), suggesting that this decline occurred in parallel for both groups. To further characterize how these disruptions in feedback sensitivity manifest at the computational level, we fit reinforcement learning models to quantify changes in the underlying parameters that govern learning and choice behavior.

### Reinforcement Learning Models Reveal Altered Learning Dynamics in NMDAR Ablation Animals

#### RL-Choice Kernel Model Accurately Captures Choice Dynamics

To characterize the computational processes underlying performance in the restless bandit task, we compared standard Q-learning with reinforcement-learning models that incorporated a choice kernel (RLCK; **Figure 2A**). The RLCK model provided the best account of behavior, yielding the lowest mean AIC (283.1) and BIC (297.9) across animals, and the highest one-step prediction accuracy (median 85 percent in LacZ and 89 percent in Grin1; **Figure 2D**). The RLCK variant with asymmetric learning rates performed similarly (ΔAIC ≈ 0 ± 3.7; ΔBIC ≈ 3.7 ± 3.8) but did not offer consistent advantages, whereas all other models showed substantially poorer fits (ΔAIC ≥ 12; ΔBIC ≥ 8; **Figure 2D**). Given its parsimony and strong predictive performance, the additive-gains RLCK model was selected for all subsequent analyses.

**Figure 2.**
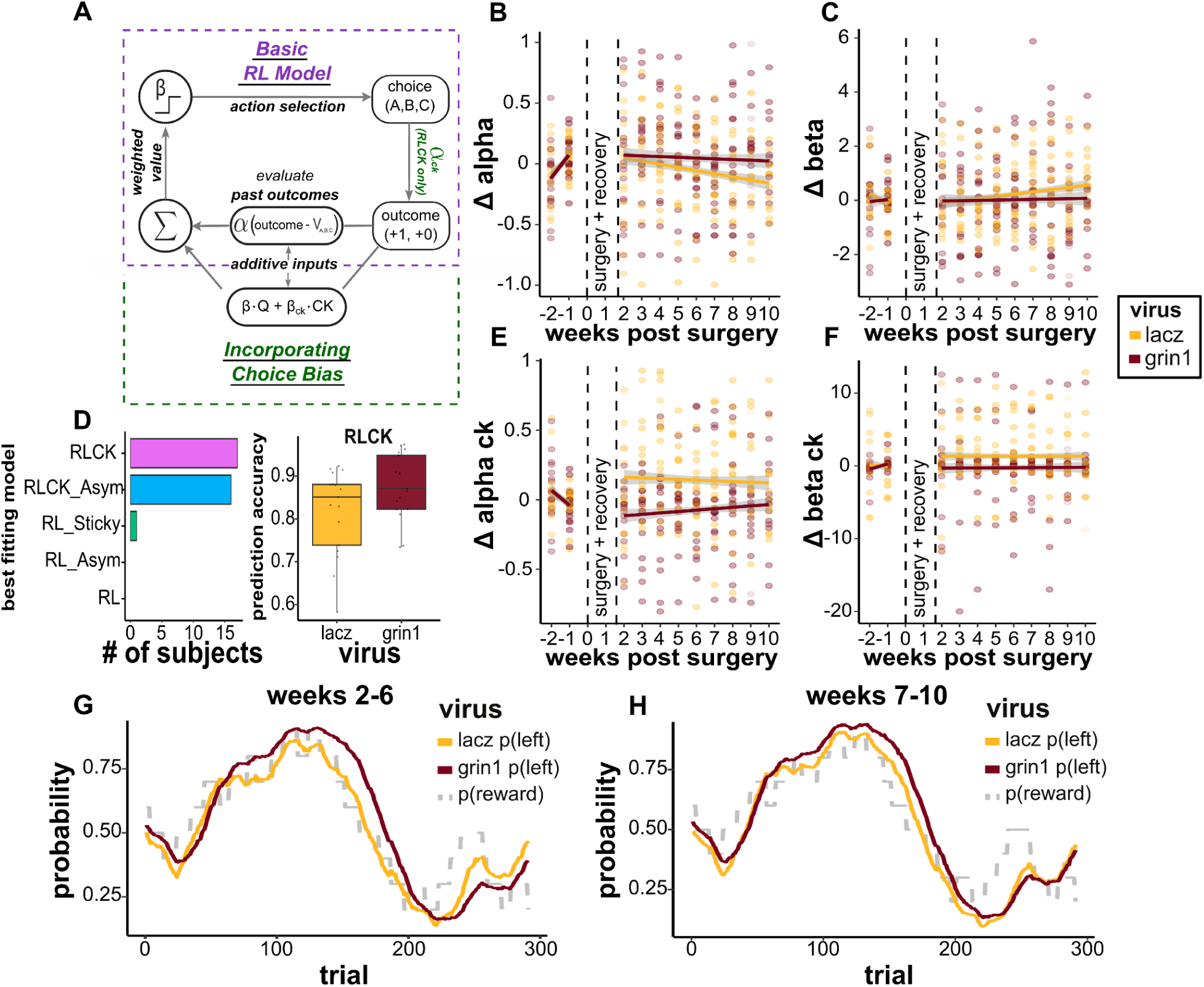
Prefrontal NMDAR ablation alters reinforcement learning dynamics in the restless bandit task. (**A**) Schematic of reinforcement learning (RL) and reinforcement learning with choice kernel (RLCK) models. (**B**) Change in learning rate (Δα) across weeks. No reliable effects were observed. (**C**) Change in inverse temperature (Δβ) across weeks. No significant main or interaction effects. (**D**) Model comparison across RL, RLCK, RLCK + Asym, RL_Asym, and RL_Sticky models based on AIC and BIC. Bars show the number of animals best fit by each model, with RLCK yielding the lowest mean AIC and BIC across animals. The asymmetric RLCK variant provided a comparable fit, while all other models showed higher information-criterion penalties. The RLCK model also achieved high one-trial-ahead prediction accuracy. (**E**) Change in choice-kernel learning rate (Δα_CK) across weeks. LacZ animals showed higher Δα_CK than Grin1, with no effect of week. (**F**) Change in inverse temperature of the choice kernel (Δβ_CK) across weeks. No reliable effects were detected. (**G**) Simulated choice probabilities from weeks 2–6 using fitted RLCK parameters over an example outcome stream (gray bar = reward probability of the left option). (**H**) Simulated choice probabilities from weeks 7–10 using fitted RLCK parameters over the same outcome stream.

#### NMDAR Ablation Selectively Alters Learning and Choice-History Dynamics

We next examined how model parameters changed from pre-to post-surgery. Learning rate (Δα), inverse temperature (Δβ), and the choice-kernel inverse temperature (Δβ_CK) showed no group differences (all p > 0.18), indicating that basic value updating and value sensitivity were not differentially impacted by virus (**Figures 2B, 2C, 2F**). In contrast, a clear group effect emerged in the choice-kernel learning rate (Δα_CK), which captures the tendency to repeat a response regardless of reward. LacZ animals showed higher Δα_CK values than Grin1 animals (β = 0.29, p = 0.008), consistent with a gradual strengthening of stable, history-based tendencies across training. Grin1 animals showed lower and more variable Δα_CK values, reflecting weaker integration of recent choices and a reduced ability to transition toward efficient, habit-like strategies (**Figure 2E**). No effects of week or virus × week interaction were observed (both p > 0.18). Together, these results indicate that prefrontal NMDAR ablation selectively disrupts the adaptive balance between value-based updating and experience-driven choice persistence.

To illustrate how these parameter differences translated to observable behavior, we simulated agents using group-level estimates from early and late post-surgical weeks under identical reward sequences. Grin1 agents consistently showed impaired tracking of dynamic reward probabilities across time, mirroring the persistent behavioral deficits observed experimentally. LacZ agents, by contrast, exhibited broader and less precise decision patterns in later weeks, consistent with an increasing reliance on a habitual and computationally efficient strategy both soon after Grin1 ablation and many weeks afterwards (**Figure 2G, 2H**). These simulations reinforce the model-based findings by providing an intuitive visualization of how parameter changes relate to patterns of adaptive versus inflexible choice behavior.

### Systemic NMDAR Antagonism Further Reduces Performance in NMDAR Ablation Animals

Administration of MK-801 markedly reduced value discrimination (p(best choice)) in both virus groups, indicating broad performance impairment under systemic NMDAR blockade. LacZ animals outperformed Grin1 under no-injection conditions (Wilcoxon rank-sum test, p < 0.001), showed a nonsignificant trend under saline (Wilcoxon rank-sum test, p = 0.068), and did not differ under MK-801 (Wilcoxon rank-sum test, p = 0.74). MK-801 reduced accuracy relative to saline within both groups (LacZ Wilcoxon signed-rank test, p = 0.002; Wilcoxon signed-rank test, p = 0.002).(**Figure 3A**). This pattern suggests partial occlusion, consistent with overlapping mechanisms of disruption. Session-level trajectories reveal that LacZ animals showing decrements on MK-801 days that rebounded following saline administration, while Grin1 animals displayed more impaired performance across all treatments, although with evidence of further impairment by MK-801 (β(MK-801) = −0.090 ± 0.014, t(278) = −6.55, p < 0.001; β(virus) = −0.029 ± 0.015, t(76) = −2.01, p = 0.048; interaction n.s.) (**Figure 3B**). Under control conditions, LacZ animals outperformed Grin1 (no-injection: Wilcoxon rank-sum test, p < 0.001; saline: Wilcoxon rank-sum test, p = 0.068), whereas under MK-801 the groups did not differ (Wilcoxon rank-sum test, p = 0.74). These findings show that systemic blockade disrupts value discrimination broadly, but with a smaller additive effect when prefrontal NMDAR signaling is already reduced.

**Figure 3.**
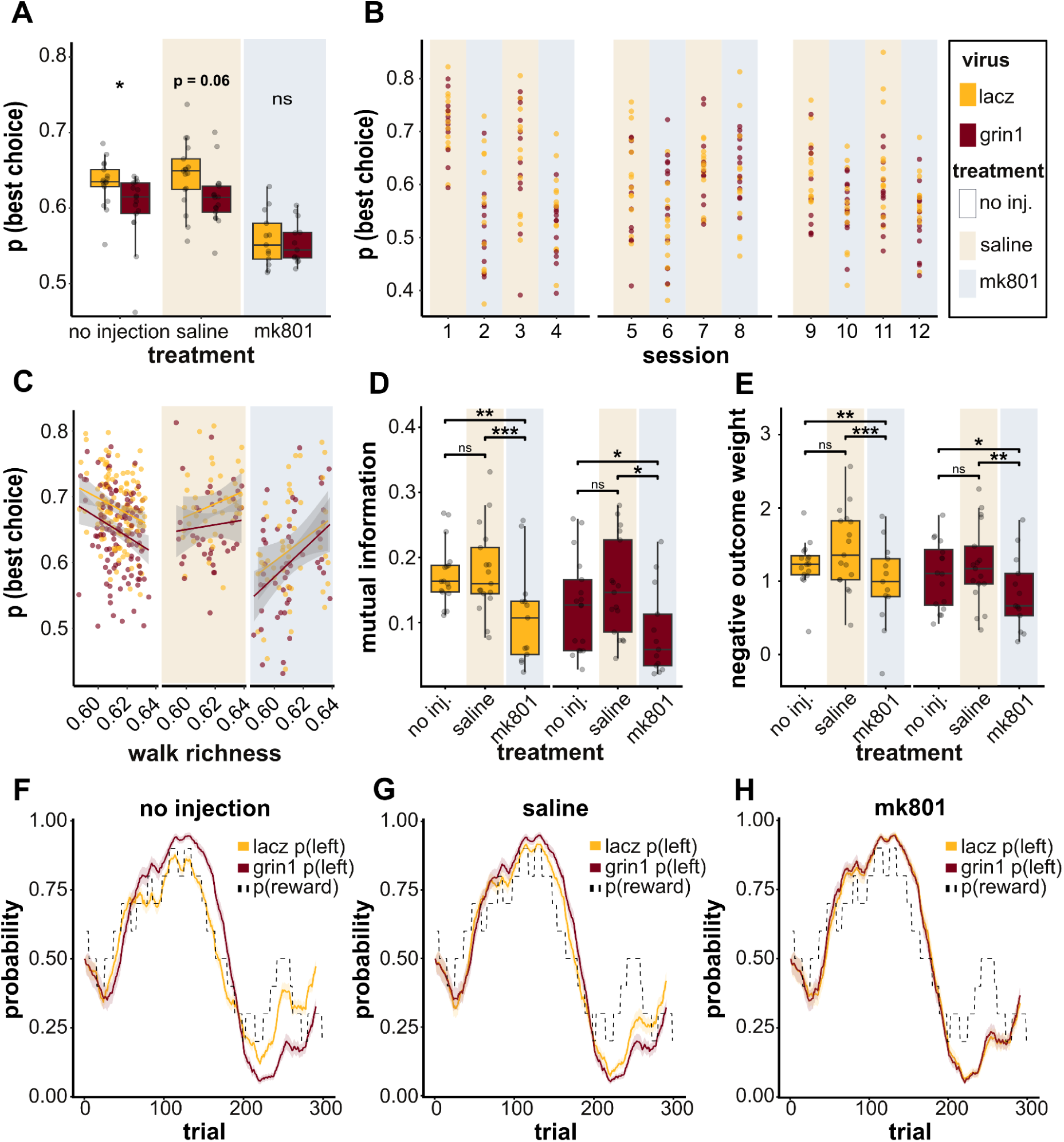
Systemic NMDA antagonism and targeted prefrontal NMDAR ablation yield overlapping yet dissociable effects in the restless bandit task. (**A**) *p*(best choice) compared between virus groups within each treatment. (**B**) Session-by-session p(best choice) across the pharmacology phase. Gaps in between sessions were not tested (eg, weekends). (**C**) Correlation between p(best choice) and environmental richness by virus within each treatment. (**D**) Mutual information (MI), reflecting the dependence of the current choice on the previous trial’s choice-outcome state (left/right × reward/no reward), compared across treatments within each virus group. (**E**) Negative outcome weight compared across treatments within each virus group. (**F**) Simulated choice probabilities under no-injection conditions using mean RLCK parameters over an example outcome stream (gray bar = reward probability of the left option). (**G**) Simulated choice probabilities under saline conditions using mean RLCK parameters over the same outcome stream. (**H**) Simulated choice probabilities under MK-801 conditions using mean RLCK parameters over the same outcome stream.

### Richness-Accuracy Relationships Diverge Across Pharmacological Conditions

We next examined how value discrimination related to walk richness across treatments. To ensure comparability across conditions, we restricted the analysis to the shared richness range observed during saline and MK-801 sessions. Within this range, clear group differences persisted under no-injection conditions, with accuracy declining as richness increased (LacZ ρ = −0.09 ± 0.09, t(12) = −1.01, p = 0.33; Grin1 ρ = −0.25 ± 0.09, t(12) = −2.90, p = 0.013). The between-group difference in correlation strength (slope) was nonsignificant (Wilcoxon rank-sum test, p = 0.24). Under saline, this relationship was attenuated and no longer significant for either group (LacZ ρ = 0.12 ± 0.20, p = 0.57; Grin1 ρ ≈ 0.00 ± 0.19, p = 1.00). The LacZ–Grin1 comparison was again nonsignificant (Wilcoxon rank-sum test, p = 0.62). In contrast, MK-801 reversed the pattern: both groups trended toward better discrimination in richer environments, reaching significance in Grin1 animals (ρ = 0.46 ± 0.14, t(12) = 3.43, p = 0.005) and remaining nonsignificant in LacZ (ρ = 0.17 ± 0.17, p = 0.33). The between-group comparison was also nonsignificant (Wilcoxon rank-sum test, p = 0.25). This shift suggests that systemic antagonism alters how reward information is retained and integrated, producing learning dynamics distinct from targeted prefrontal NMDAR disruption (**Figure 3C**).

### MK-801 Suppresses Reward History Integration and Negative Feedback Sensitivity

Because variance was higher in the saline condition (reflecting fewer sessions than no injection), virus group differences were less reliable for these measures. We therefore focused on within-group comparisons across treatments. To capture the broader integrative effects of systemic antagonism, we included mutual information (MI), which indexes how much the previous trial’s choice-outcome combination (L/R × reward/no reward) predicts the current choice, providing a compact measure of dependence on recent experience. While this metric was not emphasized in the non-pharmacological analyses, it is well suited for assessing the widespread cognitive impact of MK-801. MI analyses showed no significant difference between no-injection and saline conditions within either group (LacZ Wilcoxon signed-rank test, p = 0.53; Grin1 Wilcoxon signed-rank test, p = 0.24). In contrast, MI decreased significantly under MK-801 relative to saline in both groups (LacZ Wilcoxon signed-rank test, p = 0.002; Grin1 Wilcoxon signed-rank test, p = 0.018), indicating a broad disruption of reward-history integration (**Figure 3D**). Negative outcome-weight analyses showed a comparable pattern. Outcome weight did not differ between no-injection and saline conditions for either virus group (Wilcoxon signed-rank tests, LacZ p = 0.094; Grin1 p = 0.124), but both groups showed significant decreases under MK-801 relative to saline (Wilcoxon signed-rank tests, LacZ *p* = 0.002; Grin1 *p* = 0.004) (**Figure 3E**). These reductions suggest that systemic NMDA receptor blockade may diminish the use of negative feedback to guide future choices, consistent with impaired value-based learning and a weakened capacity to adjust behavior following unrewarded outcomes.

### Simulations Reveal Converging Impairments Under Systemic Antagonism

To link RLCK parameter changes to observed behavior, we simulated LacZ and Grin1 agents within each treatment using group-level mean parameters and shared outcome streams. Successful agents/animals would be expected to show choice probabilities that closely track actual reward probability, as represented by the dashed lines in **Figure 3F-H**. No-injection simulations replicated stable deficits in Grin1 agents’ ability to track shifting reward probabilities relative to LacZ(**Figure 3F**). Under saline, simulated LacZ agents showed reduced adaptability, narrowing the gap between groups and suggesting a mild shift toward slower, more habitual strategies (**Figure 3G**). Under MK-801, simulated trajectories for both groups less closely tracked reward probability, and group behavior became nearly indistinguishable (**Figure 3H**). These results indicate that systemic NMDAR blockade broadly suppresses adaptive learning, collapsing previously distinct computational profiles into a shared impaired state. The attenuated difference between groups under MK-801 supports the interpretation that prefrontal NMDAR ablation occludes much of the additional disruption produced by systemic antagonism.

## Discussion

We used a CRISPR-Cas9 viral strategy to selectively ablate the GluN1 subunit of NMDAR in the mPFC and benchmarked its effects against systemic blockade with the NMDAR antagonist MK-801. This targeted model produced enduring impairments in decision making that reflect core cognitive symptoms of schizophrenia, including reduced value discrimination, altered feedback sensitivity, and difficulty adjusting to changing contingencies[6,30]. Systemic antagonism produced broader impairments across both groups of animals, reducing accuracy, mutual information, and negative outcome weight regardless of viral condition. The attenuated effects of MK-801 in Grin1 animals imply that once prefrontal NMDAR signaling is removed, systemic antagonism has limited additional influence. However, these limited impacts of NMDAR antagonism on ablation animals may suggest either incomplete ablation in mPFC and/or recruitment of hippocampal and striatal networks that support learning and flexibility[43,44]. Taken together, these findings show that local and systemic manipulations interact but are not interchangeable, providing complementary views on how glutamatergic dysfunction contributes to schizophrenia-related decision making deficits [17,22].

Reinforcement-learning analyses further clarified the computational mechanisms underlying these behavioral patterns. The RLCK model, which incorporates a choice kernel alongside value updating, consistently provided the best account of choice behavior in both groups, outperforming standard Q-learning and asymmetric or sticky-choice variants. This strong performance indicates that recent choice history is a central component of policy formation in this task. Impaired animals retained enough value tracking to perform above chance, suggesting that behavior reflects an interaction between value-based updating and choice persistence rather than simple repetition. Adding asymmetric learning rates did not improve fits, pointing to group and treatment differences arising from the balance between value-based updating and choice-history influences. These results align with behavioral evidence of reduced negative feedback weighting and indicate that value updating and choice persistence together characterize how NMDAR disruption alters decision making.

Our findings are consistent with the idea that NMDAR signaling in the prefrontal cortex supports credit assignment, the process of linking actions to delayed outcomes during learning[45]. Disruption of this mechanism likely forces animals to rely more on recent history signals, reflected in stronger influence of the choice kernel. In computational terms, prefrontal NMDAR hypofunction may reduce the precision of prediction-error weighting, producing shallow updating and weaker discrimination between informative and uninformative outcomes[22]. Similar mechanisms appear in hierarchical learning and predictive coding accounts of schizophrenia, where diminished NMDAR-mediated plasticity weakens flexible reweighting of action-outcome relationships. At a systems level, these cognitive changes likely reflect altered interactions within prefrontal-striatal networks that arbitrate between goal-directed and habitual control[44]. NMDAR-dependent signaling in mPFC shapes striatal plasticity and reinforcement sensitivity, suggesting that its loss biases animals toward striatal-driven, perseverative strategies[46,47].

In our task, control animals gradually shifted toward a more habitual strategy as they accumulated experience, whereas Grin1 animals appeared to enter this mode prematurely, relying on simplified choice rules even when flexibility remained beneficial. Although performance deficits in Grin1 animals persisted throughout the post-surgical period, the trajectory in Figure 1D suggests a modest trend toward partial behavioral recovery over time. This trend did not reach statistical significance, but it does raise the possibility that some degree of functional compensation may occur despite ongoing receptor loss. This interpretation aligns with prior work in this mouse model showing that NMDAR ablation triggers compensatory increases in both excitatory and inhibitory signaling within mPFC[33]. Such synaptic rebalancing could help stabilize behavior and prevent further decline, even though it is insufficient to restore full value-based learning and decision-making capacity.

These behavioral and computational patterns parallel findings in schizophrenia, where impaired goal-directed updating coexists with exaggerated response persistence[2,48]. Reduced negative-feedback sensitivity, stable low learning rates, and elevated choice persistence in our ablation animals mirror core computational phenotypes identified in human reinforcement learning studies. Integrating trial-level modeling with circuit-level measures will help clarify how prefrontal NMDAR signaling shapes precision weighting, feedback-driven learning, and transitions between flexible and habitual control. By linking mechanistic accounts of synaptic plasticity with quantitative behavioral signatures, this work strengthens the bridge between animal and human studies and supports future interventions targeting glutamatergic modulation.

## Acknowledgements

This work was supported by NIH P50MH119569 and NIH R01MH123661.

---

**Supplementary Figure 1.**
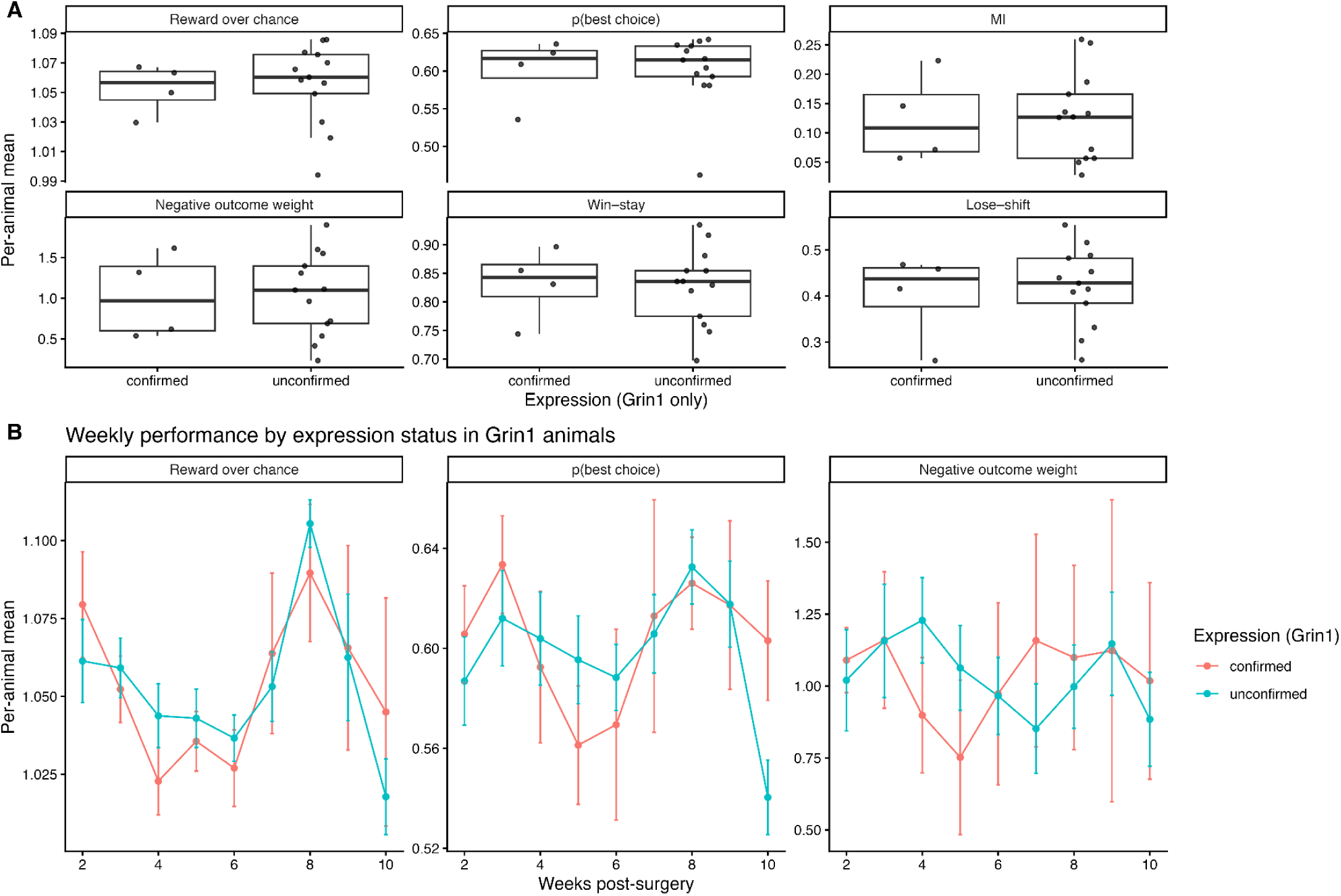
Behavioral effects of confirmed versus unconfirmed Grin1 expression. **(A)** Core behavioral measures for Grin1 animals with confirmed and unconfirmed expression across post-surgery no-injection sessions. Each point reflects a per-animal mean, plotted for reward over chance, p(best choice), mutual information (MI), negative outcome weight, win–stay, lose–shift, and the correlation between p(best choice) and walk richness (ρ). None of these measures differed significantly between expression groups (Wilcoxon rank-sum tests, all p > 0.61; n = 4 confirmed, n = 13 unconfirmed). **(B)** Weekly trajectories of reward over chance, p(best choice), and negative outcome weight across weeks 1–10 post-surgery in confirmed and unconfirmed Grin1 animals. Linear mixed-effects models showed no main effect of expression on reward over chance (p = 0.71), p(best choice) (p = 0.80), or negative outcome weight (p = 0.55), and no significant expression × week interactions (all p > 0.15). These results indicate that confirmed and unconfirmed Grin1 expression groups exhibit comparable behavioral profiles across all measures relevant to the main text.

## Supplemental Methods

### Behavioral Training

#### Pre-Training Familiarization

To ensure familiarity with the liquid reinforcer, mice were given overnight access to 50 % water-diluted vanilla Ensure in their home cage. Upon reaching appropriate age for operant training, mice were placed in the operant touchscreen chamber for a 30-minute habituation session. A single 50 µL aliquot of reward was dispensed at the start of the session, allowing animals to freely explore the chamber and retrieve the reward in the absence of operant demands.

#### Touchscreen Acquisition Stages

##### Stage 1 – Initial Touch Training

In this introductory phase, mice were trained to associate screen contact with reward delivery. A 7 µL reward was dispensed automatically every 30 seconds regardless of behavior. In addition, if a mouse touched any portion of the screen within the region displaying a single stimulus image, it received an additional reward of 21 µL. Sessions lasted 30 minutes, and mice advanced to the next stage once they completed ≥30 trials within session on two consecutive days.

##### Stage 2 – Must Touch Training

During this phase, the time-based reward delivery was eliminated. Mice were required to make a screen touch within the image region to receive a 7 µL reward. Session duration remained 30 minutes. Advancement criteria were identical to Stage 1: 30 completed trials in 30 minutes for two consecutive sessions.

##### Stage 3 – Must Initiate Training

This stage introduced a requirement to self-initiate trials. After completing a rewarded response and a 3-second inter-trial interval (ITI), a light cue at the reward port signaled that the next trial could begin. The mouse initiated the trial by entering the port, triggering stimulus presentation on the screen. A correct screen touch led to delivery of a 7 µL reward. Sessions were 30 minutes long, and mice progressed once they completed ≥30 trials in 30 minutes across two sessions.

##### Stage 4 – Punish Incorrect Training

This schedule introduced a penalty for incorrect responses. Mice were presented with a single illuminated image on either side of the screen and were required to respond selectively. Touches to blank screen areas (non-image side) resulted in a 10-second timeout, accompanied by a flashing house light. Correct responses triggered a 7 µL reward, followed by a 3-second ITI and trial re-initiation. Sessions continued for up to 60 minutes or until 200 trials were completed. Advancement required 200 trials per session on two consecutive days.

##### Stage 5 – Deterministic Image Discrimination (100–0) Training

This stage marked the transition to value-based decision-making. Mice were presented with two images simultaneously, one in each screen aperture. One image was always rewarded (100%), while the other was never rewarded (0%), independent of location. No punishment was delivered for incorrect choices. Sessions lasted 120 minutes or until 250 trials were completed. Mice were advanced when they consistently completed 250 trials with ≥85% accuracy.

##### Stage 6 – Probabilistic Spatial Discrimination Training

This training block introduced stochastic and reversible reward contingencies. Trials displayed identical stimuli on the left and right side of the screen, and mice had to learn which side offered a higher probability of reward. The task began with a 90/10 probability schedule, followed by an 80/20 schedule, and then two days of 70/30 probability sessions. Contingencies reversed once the mouse selected the high-probability option on 9 out of the previous 10 trials. Each session concluded after 300 trials or 120 minutes. This training prepared animals for dynamic reward environments encountered in the bandit task.

### Histology and viral injection localization

#### Perfusion and tissue processing

Following the final behavioral session, mice were deeply anesthetized with Euthaphen and transcardially perfused with phosphate-buffered saline (PBS), followed by 4% paraformaldehyde (PFA) in PBS. Brains were extracted, post-fixed in 4% PFA for 24 h at 4 °C, then transferred to PBS containing sodium azide for storage. Prior to sectioning, tissue was cryoprotected in 15% and then 30% sucrose in PBS (with sodium azide) until sinking. Coronal sections (30 µm) spanning the targeted medial prefrontal cortex were cut on a cryostat and moved to 12 well plates for staining.

#### Immunofluorescence

Sections were rinsed in PBS and blocked for 1 h at room temperature in PBS with 0.3% Triton X-100 and 5% normal goat serum. Slices were incubated overnight at 4 °C with rat anti-mCherry monoclonal antibody (clone 16D7; ThermoFisher, Cat. #M11217; typical dilution 1:500 in blocking buffer). The following day, sections were washed in PBS and incubated for 2 h at room temperature with goat anti-rat IgG (H+L) Alexa Fluor 594 (ThermoFisher, Cat. #A-11007; typical dilution 1:1000). Sections were coverslipped with an anti-fade mounting medium. All steps used light protection to preserve fluorophore signal. Reagent sources and catalog numbers are reported above, blocking serum species matched the secondary antibody host.

#### Imaging and localization

Fluorescent images were acquired on a Zeiss Axiolab 5 microscope using identical exposure settings across animals within each staining batch. Injection sites and spread were identified by mCherry immunofluorescence and native reporter signal. Localization was confirmed by comparison to a mouse brain atlas [49]. During the staining and imaging process we ran into issues with tissue storage, resulting in degradation of tissue quality that prevented proper imaging in some of the Grin1 animals. To assess potential confounds arising from this, animals with confirmed versus unconfirmed expression were compared across all behavioral analyses presented in this paper (**Supplemental Figure 1**). No significant differences were found in any behavioral metric, indicating that inclusion of unconfirmed cases is unlikely to have biased the reported findings.

### Behavioral Apparatus

All behavioral training and testing were conducted in Bussey–Saksida touchscreen operant chambers (Lafayette Instrument Company, Lafayette, IN) housed in sound-attenuating cubicles. Chambers featured black Plexiglas walls, a transparent lid, a stainless steel grid floor, and an infrared touchscreen that registered both discrete and repeated touches with high spatial resolution. A standard mask restricted responding to two apertures on the left and right sides of the screen, which displayed the stimuli used throughout training and testing and were positioned to allow direct nosepoke responses. Liquid reinforcement (50% water-diluted vanilla Ensure) was delivered in calibrated volumes through a recessed port opposite the touchscreen, equipped with infrared sensors to detect reward collection. All task events, including stimulus presentation, response registration, reward delivery, and timing, were controlled and logged using ABET II software.

### Data Analysis

#### Mutual information

We quantified the dependence of current choice on recent history with mutual information. For each animal we computed: I(choice_t; [choice_{t-1}, reward_{t-1}]) and, for comparison, I(choice_t; choice_{t-1}). All variables were categorical. Values were averaged within condition.

#### Negative outcome weight

Negative outcome weight indexed how much unrewarded outcomes promoted switching relative to rewarded outcomes, normalized by the overall switching rate:

Negative outcome weight = [ P(switch | no reward) – P(switch | reward)] / P(switch). Here, “switch” means changing sides from trial t-1 to trial t. Conditional probabilities were estimated from empirical frequencies per animal and then aggregated within condition.

#### Reinforcement-learning models

We fit a series of reinforcement-learning (RL) models to trial-by-trial behavior, each capturing distinct computational mechanisms underlying choice. All models generated action probabilities via a softmax function applied to decision logits, and parameters were estimated separately for each animal and schedule using bounded L-BFGS-B optimization. Below, *a* denotes the chosen action, *ā* the unchosen action, and *r□* ∈ {0,1} the reward outcome.

#### RL (Rescorla–Wagner)

Parameters: α (learning rate), β (inverse temperature) This baseline model updates the value of the chosen action based on the reward prediction error:

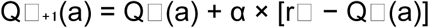

Choice probabilities are determined by a softmax transformation over action values:

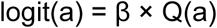

#### RLCK (RL with Choice Kernel, additive gains form)

Parameters: α, β, α_C (choice kernel update rate), β_CK (choice kernel weight)

This model augments the standard RL process with a choice kernel (CK) that captures recent choice tendencies independent of reward.

After each trial:

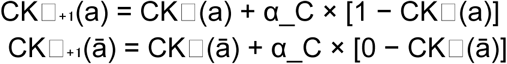

The value and choice bias signals combine additively in the decision logits:

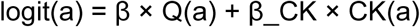

#### RLCK_Add_Asym (Additive Gains RLCK with asymmetric learning)

Parameters: α_pos, α_neg, β, α_C, β_CK

This extension allows distinct learning rates for rewarded and unrewarded outcomes, capturing asymmetric reward sensitivity:

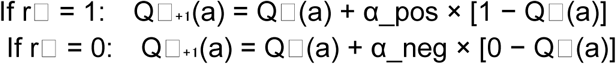

The CK updates as in the standard RLCK model, and the combined decision logits remain:

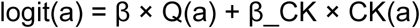

#### RL_Sticky (RL with stay bias)

Parameters: α, β, β_stay

A binary “stay” indicator *s□* (reflecting whether the previous choice was repeated) augments the decision logits:

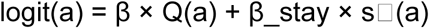

where *s□* = [1,0] if the previous choice was left, [0,1] if right, and [0,0] on the first trial of a block.

#### RL_Asym (Asymmetric learning rates)

Parameters: α_pos, α_neg, β

This model uses separate learning rates for positive and negative outcomes but omits the choice kernel term:

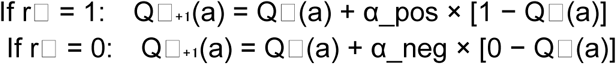

Choice probabilities are given by:

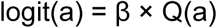

#### Estimation and model comparison

For each subject we used maximum likelihood with multi-start optimization and simple box constraints on parameters. We recorded convergence status, best-fitting parameters, and negative log-likelihood (NLL). Model evidence was summarized with:

● AIC = 2*k + 2*NLL
● BIC = k*ln(n) + 2*NLL
● Akaike weights: w_i = exp(−0.5*DeltaAIC_i) / sum_j exp(−0.5*DeltaAIC_j)

We report mean Akaike weight, median DeltaAIC, and number of subjects for which each model achieved the lowest AIC.

## Notes

### Competing Interest Statement

The authors have declared no competing interest.

